# Combined Effect of Shear Stress and Laser-Patterned Topography on Schwann cell Outgrowth: Synergistic or Antagonistic?

**DOI:** 10.1101/2020.07.09.195420

**Authors:** Eleftheria Babaliari, Paraskevi Kavatzikidou, Anna Mitraki, Yannis Papaharilaou, Anthi Ranella, Emmanuel Stratakis

## Abstract

Although the peripheral nervous system exhibits a higher rate of regeneration than that of the central nervous system through a spontaneous regeneration after injury, the functional recovery is fairly infrequent and misdirected. Thus, the development of successful methods to guide neuronal outgrowth, *in vitro*, is of great importance. In this study, a precise flow controlled microfluidic system with specific custom-designed chambers, incorporating laser-microstructured polyethylene terephthalate (PET) substrates comprising microgrooves, was fabricated to assess the combined effect of shear stress and topography on Schwann cells’ behavior. The microgrooves were positioned either parallel or perpendicular to the direction of the flow inside the chambers. Additionally, the cell culture results were combined with computational flow simulations to calculate accurately the shear stress values. Our results demonstrated that wall shear stress gradients may be acting either synergistic or antagonistic depending on the substrates groove orientation relative to the flow direction. The ability to control cell alignment *in vitro* could potentially be used in the fields of neural tissue engineering and regenerative medicine.

## 1. Introduction

Considering that the neurological injuries cannot typically self-recover, there is a need for developing strategies to stimulate neurogenesis. Furthermore, following damage in the peripheral nervous system (PNS), regeneration is usually unsuccessful since axons are directed towards inappropriate targets [1]. As a consequence, neural tissue engineering has emerged as a promising field for the development of new graft substitutes [2]. Indeed, the ultimate goal of using a tissue-engineered construct is to adequately mimic both the topographic features of the extracellular matrix (ECM) as well as its surrounding stimuli, including for example mechanical stresses and soluble factors, so that cells will respond within the artificial environment as they would *in vivo* during development.

It has been well-reported that the surface topography significantly affects the adhesion, orientation, proliferation, and differentiation of cells [3–9]. Therefore, many groups have focused their interest on the controlled modification of materials’ surfaces, as a strategy for guiding the neurite outgrowth, which is crucial for the development of functional neuronal interfaces [10–13]. Among various techniques employed, ultrashort-pulse laser structuring has proved to be important for engineering surface topography in various materials [13–22] including silicon (Si), PET, and poly(lactide-co-glycolide) (PLGA). Indeed, the laser-induced topography was found to significantly affect the neuronal adhesion, growth, and orientation.

Another crucial factor in neuronal regeneration, that should not be overlooked, is the mechanical environment provided by the ECM. Flow-induced shear stress, in particular, influences mechanoreceptors, like ion channels and focal adhesions, as well as responses, such as nitric oxide and intracellular calcium production and cytoskeletal remodeling [23]. Shear stress is applied at discrete local points and is transmitted through the cell body along cytoskeletal microstructures, which in turn trigger intracellular mechanical signaling. Thus, shear stress alters not only the cell’s shape but also the intracellular signaling pathways [1,24]. Flow-induced shear stress can be applied to cells, *in vitro*, using specially designed microfluidic systems. In such systems, laminar flow replicates the physiological fluid flow inside the body, facilitates mass transport of solutes and supplies consistent nutrient delivery and effective waste removal resulting in a more *in vivo*-like environment [25]. Indeed, the continuous flow of nutrient is a distinct advantage that microfluidics bring to dynamic cell cultures in contrast to conventional static ones.

Many studies have shown that the dynamic culture of neuronal and/or glial cells has a positive effect on their proliferation and differentiation [26–31]. Specifically, Millet et. al. [26] developed a specific culture system that showed increased viability and channel-length capacity of primary hippocampal neurons. Moreover, Chung et. al. [27] developed a PDMS gradient-generating microfluidic platform that exposed human neural stem cells (hNSCs) to a concentration gradient of known growth factors (GF), under continuous flow (5 10^-5^ Pa). As a result, a minimization of autocrine and paracrine signaling was observed. Additionally, they found that the differentiation of hNSCs into astrocytes was inversely proportional to GF concentration while proliferation was directly proportional. Gupta et. al. [28,29] used an *in vitro* model to apply shear stress on primary Schwann cells in the form of laminar fluid flow (3.1 Pa for 2 hours). They observed increased proliferation and down-regulation of two pro-myelinating proteins, myelin-associated glycoprotein (MAG) and myelin basic protein (MBP). These results implied that a low level of mechanical stimulus may directly trigger Schwann cell proliferation. Furthermore, Majumdar et. al. [30] developed a polydimethylsiloxane (PDMS) microfluidic cell co-culture platform where hippocampal neurons and glia maintained for several weeks. In particular, co-culture with glia provided nutrient media for maintaining healthy neural cultures and enhanced the transfection efficiency of neurons in the platform. Similarly, Shi et. al. [31] fabricated two PDMS microfluidic cell culture systems. They observed that the co-culture of neurons and glia increased the number and stability of synaptic contacts, as well as, the secreted levels of soluble factors. These results confirmed the importance of communication between neurons and glia for the development of stable synapses in microfluidic platforms.

However, the combined effect of flow-induced shear stress and surface topography on neurite outgrowth has been rarely reported. Specifically, Kim et. al. [1] studied the effect of mechanical stimulation on PC12 cells cultured in microfiber-based substrates. They observed that the shear stress affected the length and orientation of neurons along the microfibers. Jeon et. al. [32] investigated the combined effects of surface topography and flow-induced shear stress on the neuronal differentiation of human mesenchymal stem cells. They applied different shear stresses in a PDMS substrate with micrometric grooves. An increased directionality under flow conditions was observed. Moreover, an increased neurite length was noticed on the seventh day. However, a significant decrease in neurite length was observed on the tenth day. These preliminary results were not conclusive on how the combined effect of shear stress and topography affects neuronal growth. It should also be emphasized that in both studies, the flow was not continuous but was rather applied for only a few hours per culture day. As a result, the *in vivo* dynamic culture conditions may not be adequately simulated *in vitro.*

The present work aims to present a first study of the combined effect of shear stress and topography on the adhesion, growth, and orientation of Schwann cells under dynamic culture conditions attained via continuous flow. Schwann cells are important glial cells of the PNS, providing both molecular and topographical guidance stimuli for the development and outgrowth of neurons [33]. For this purpose, we developed a precise flow controlled microfluidic system with a custom-designed chamber incorporating laser-microstructured PET culture substrates comprising microgrooves (MG) [18]. PET has been widely used for cell culturing, surgical suture material and prosthetic vascular grafts due to its biocompatibility and its excellent mechanical strength and resistance [34,35]. The MG were positioned either parallel or perpendicular to the direction of the flow and the response of Schwann cells was evaluated in terms of adhesion, orientation, and cell length. The cell culture results were combined with computational flow simulation studies employed to accurately calculate the shear stress values. Our findings demonstrated the ability to guide the outgrowth of Schwann cells, *in vitro*, via flow-induced shear stress and surface topography, which is crucial for neural tissue regeneration.

## 2. Materials and Methods

### 2.1 Design of the Microfluidic System

The microfluidic system (Figure 1a) is composed of an air compressor (Durr Technik, USA) and an OB1 pressure controller (Elveflow, France) which are connected, through silicon tubing, to the nutrient reservoirs (Elveflow, France). Then nutrient (in this case culture medium), through poly(tetrafluoroethylene) (PTFE) tubing (interior diameter 0.5 mm), moves to the flow sensors (Elveflow, France), which precisely control the flow rates, to the bubble traps (Elveflow, France), to the custom-made geometry chambers (Ebers, Spain) containing the laser-microstructured substrates with the cells and finally to the waste reservoirs (Elveflow, France). The laser-microstructured substrates were placed parallel or perpendicular to the direction of the flow inside the chambers. The chambers consist of polysulfone upper lids for loading cells (length: 6 mm, width: 6 mm, height: 13 mm) and for flowing fluids across the cells and the laser-microstructured substrates (length: 11 mm, width: 6 mm, height: 150 μm). The experiments were performed under continuous flow conditions. The chambers, as well as the waste reservoirs, were placed inside a 5% carbon dioxide (CO_2_) incubator at 33°C, for the whole duration of the dynamic culture experiments. The cross-section image of the chamber, containing the laser-microstructured substrates with the cells, where the flow occurs is also illustrated (Figure 1b).

**Figure 1:**
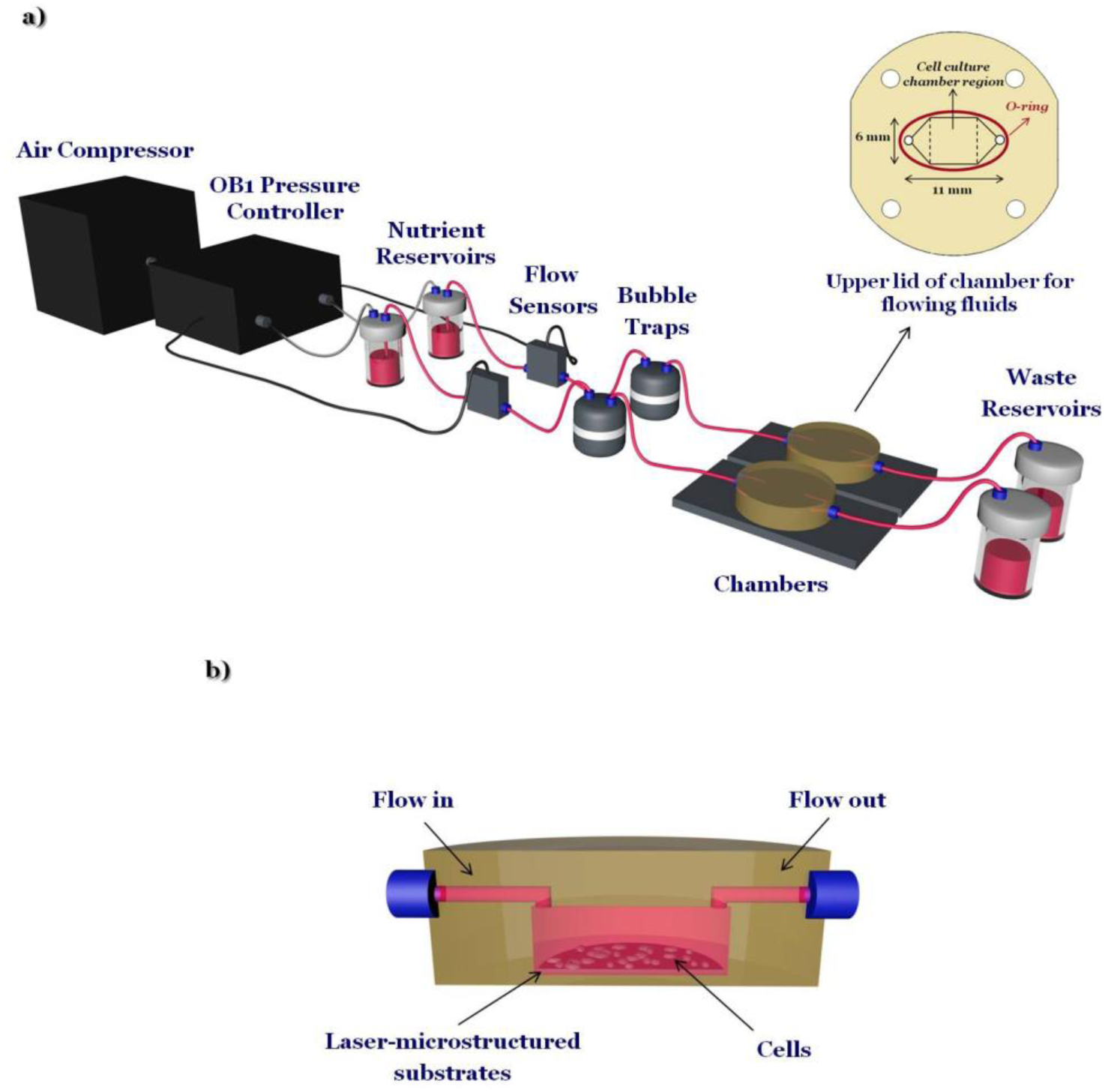
(a) Schematic illustration of the custom-designed microfluidic system. It is composed of an air compressor and an OB1 pressure controller connected with nutrient reservoirs, flow sensors, bubble traps, a couple of chambers including the cells and the laser-microstructured substrates and waste reservoirs. The upper lid of the chamber for flowing fluids across the cells and the laser-microstructured substrates is shown at the inset. Specifically, the length of the chamber’s upper lid for flow is 11 mm and the width is 6 mm. The cell culture chamber region is pointed out in the dashed lines. (b) Cross-section image of the chamber, containing the laser-microstructured substrates and cells, where the flow occurs.

### 2.2 Fabrication of Laser-Microstructured Substrates

The microstructured substrates were fabricated via ultrafast laser direct writing of PET coverslips used for cell cultures (Sarstedt, Numbrecht, Germany). This is a simple, low cost, and effective method to control the substrate topography [36]. For this purpose, a Yb:KGW laser source was used with a wavelength of 1026 nm, 1 kHz repetition rate and 170 fs pulse duration. The laser-microstructured substrates were fabricated at a constant fluence of 11.9 J/cm^2^ and scan velocity of 7 mm/s. The total patterned area was 3 mm × 3 mm while the MG periodicity was equal to 28.68 μm.

### 2.3 Morphological Characterization of Laser-Microstructured Substrates by Scanning Electron Microscopy (SEM)

Following the laser structuring process, the laser-microstructured substrates were sputter-coated with a 15 nm layer of gold (Baltec SCD 050, BAL-TEC AG, Balzers, Liechtenstein) and observed under a scanning electron microscope (JEOL JSM-6390 LV, Jeol USA Inc, Peabody, MA, USA) using an acceleration voltage of 15 kV. An image processing software, Fiji ImageJ, was used to analyze the geometrical characteristics of the MG, as described in [18]. The aspect ratio of the MG, A, was calculated to be the ratio of the MG depth (d) to their width (w) (*A* = *d*/*w*). The roughness ratio, r, was calculated to be the ratio of the actual, unfolded, surface area of MG to the total irradiated area [*r* = 1 + (2d/*w*)]

### 2.4 Static and Dynamic Cultures

Mouse Schwann cell line (SW10), an established adherent neuronal cell line, was obtained from ATCC® (Code: CRL-2766™). SW10 cells were grown in cell culture flasks using culture medium [Dulbecco’s Modified Eagle’s Medium (DMEM) (Invitrogen, Grand Island, NY, USA) supplemented with 10% Fetal Bovine Serum (FBS) (Biosera, Sussex, UK) and 1% antibiotic solution (Gibco, Invitrogen, Kalsruhe, Germany)] in a 5% CO_2_ incubator (Thermo Scientific, OH, USA) at 33°C.

Prior to any experiment, the laser-microstructured substrates, the reservoirs, the bubble traps, and the chambers were UV-sterilized. Subsequently, the substrates were transferred into sterile wells of 24-well plates (Sarstedt, Numbrecht, Germany) for static cultures or/and inside the chambers of the microfluidic system for dynamic cultures. Planar PET coverslips for cell cultures (Sarstedt, Numbrecht, Germany), termed as PET-Flat, were used as the control samples.

Prior to any culture, the substrates were coated with a 15 μg/mL solution of laminin (Sigma-Aldrich, St. Louis, MO, USA) to enhance cell adhesion. A number of 2.5 × 10^4^ cells were used for both the static and dynamic culture experiments. For the dynamic cultures, cells were seeded on the planar PET coverslips or the laser-microstructured substrates, placed inside the chamber and kept at rest overnight in a 5% CO_2_ incubator at 33°C. The next day, the continuous perfusion was initiated and lasted for 2 days. Based on the results of our previous work [18], under static conditions, we determined that it would be more effective to quantify the cell length after a 3-day culture period. Moreover, parametric studies were performed both under static and dynamic conditions before selecting these optimal parameters (dimensions of the substrate, topography of the substrate, cell densities, flow rates) for investigating the effect of shear stress (section A in the Supplementary Material).

To our knowledge, there are no available data on the shear stress values applied *in vivo* to the PNS cells. Additionally, although it is known [37] that there are small arteries, coined as *vasa nervorum*, providing blood supply to peripheral nerves, the velocity’s values are unknown. Thus, parametric studies were performed, as aforementioned, to conclude to the flow rates of 50 and 200 μL/min. We observed that at higher flow rate values cells detached from the substrates.

The mean velocity, *ū*, in the microfluidic system is estimated by the equation [38]:

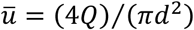

where *Q* is the flow rate and *d* is the tubing diameter (*d* = 0.5 mm).

While, the shear stress, σ, exerted on the cell layer can be estimated, in a simplistic approach, by the equation [39]:

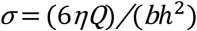

where η is the nutrient’s viscosity (*η* ∼ 0.01 gcm^-1^s^-1^ [40,41]), *Q* is the flow rate, *b* is the width of the chamber’s upper lid for flowing fluids (*b* = 6 mm) and *h* is the height of the chamber’s upper lid for flowing fluids (*h* = 150 μm).

Table 1 shows the flow rates as well as the corresponding mean velocity and shear stresses in the microfluidic system.

**Table 1:** *Values for flow rate (Q), mean velocity [ū* = (4*Q*)/(*πd*^2^)*] and shear stress [σ* = (6*ηQ)/(bh*^2^*)] in the microfluidic system* [38,39].

| $Q (\mu\text{L/min})$ | $\bar{u} (\text{m/s})$ | $\sigma (\text{Pa})$ |
| --- | --- | --- |
| 50 | 0.0042 | 0.04 |
| 200 | 0.0170 | 0.16 |

### 2.5 Computational Flow Simulations in the Microfluidic System Chamber

The commercial software ICEM-CFD v12.1 (Ansys Inc.) was used to generate the mesh of the computational model. The computational grid for the flat substrate case contains 435K hexahedral elements and non-uniform grid node spacing to produce higher grid density at the inlet and outlet regions of the square-shaped cell culture region of the chamber (Figure 1a). Near wall grid refinement was imposed through a viscous layer adjacent to the walls in order to capture the velocity gradients in the boundary layer and resolve the wall shear stress. Straight tube extensions were added to the inlet and outlet of the microfluidic system chamber to prevent upstream and downstream contamination respectively of the chamber flow domain from flow disturbances caused by forced outflow boundary conditions or non-fully developed flow entering the chamber.

For the microstructured substrate case the 3D computational grid contains 1.5M hexahedral elements and non-uniform node spacing to produce near wall refinement and higher grid density within the MG that runs parallel to the mean flow direction. In the case where the MGs are placed perpendicular to the mean flow direction symmetry of the geometry and periodicity of the MG is exploited to construct a 2D computational grid with 240K elements. In both cases it is assumed that flow is fully developed at the upstream boundary of the MG region.

The Navier–Stokes and continuity equations for incompressible steady laminar flow in the absence of body forces were solved using Fluent v12.1 (Ansys Inc.). For the flow field computations, the wall was assumed rigid and the fluid was modeled as incompressible Newtonian with a density of 1.0 gr/cm^3^ and a viscosity of 1.0 cP. A uniform velocity was applied perpendicular to the inlet boundary and a traction free boundary condition was applied at the outlet of the domain.

### 2.6 Immunocytochemical Assays

The F-actin of the cytoskeleton and the double-stranded helical DNA of the nucleus, of SW10 cells, were stained with phalloidin and 4’,6-Diamidino-2-Phenylindole (DAPI), respectively. Actin is the major cytoskeletal protein of most cells. It is highly conserved and participates in various structural and functional roles [42]. In this case, actin exists as actin filaments (F-actin) and is stained with a specific phalloidin (Alexa Fluor® 680 Phalloidin). After 3 days of culture, the samples were fixed with 4% paraformaldehyde (PFA) for 15 minutes and permeabilized with 0.1% Triton X-100 in phosphate-buffered saline (PBS) for 5 minutes. The non-specific binding sites were blocked with 2% Bovine Serum Albumin (BSA) in PBS for 30 minutes. Then, the samples were incubated for 2 hours at room temperature with Alexa Fluor® 680 Phalloidin (Invitrogen, Thermo Fisher Scientific) (1:250 in PBS–BSA 1%) for F-actin staining. Finally, the samples were washed with PBS and put on coverslips with DAPI (Molecular Probes by Life Technologies, Carlsbad, CA, USA) for nuclei staining. Cell imaging was performed using a Leica SP8 inverted scanning confocal microscope. The objective of x40 was used. To obtain the images of cells both on the top of MG and inside the MG, the z-stack of the confocal microscope was used. To investigate changes in the directional orientation of cells’ cytoskeleton, the “Local gradient orientation” for directionality was performed using the Fiji ImageJ plug-in “Directionality” [43]. In this way, the data of the amount of cells, presented in the input image, in each direction was extracted and plotted as a polar plot. To compare the different cases, the cell population per angle was normalized with the maximum value in each case and expressed as normalized cell population. Additionally, in order to determine the cell length, Fiji ImageJ and Harmony^®^ software of Operetta High-Content Imaging System (Perkin Elmer) were used.

### 2.7 Statistical Analysis

Statistical analysis of the data was performed using post hoc Tukey HSD test. A p-value < 0.05 was considered significant. For each case investigated, a series of three different experiments have been performed.

## 3. Results

### 3.1 Morphological Characteristics of Laser-Microstructured PET Substrates

The laser-structured microgrooved substrates (PET-MG) were morphologically characterized by scanning electron microscopy (SEM). The corresponding top-view and cross-section SEM images are presented in Figure 2, showing that the width, *w*, of the MG was 28.68 ± 0.47 μm, the depth, *d*, 8.87 ± 0.44 μm, the aspect ratio, *A* = *d/w* = 0.309 ± 0.01 and the roughness ratio, *r* = 1 + (2*d/w*) = 1.62 ± 0.02 [18].

**Figure 2:**
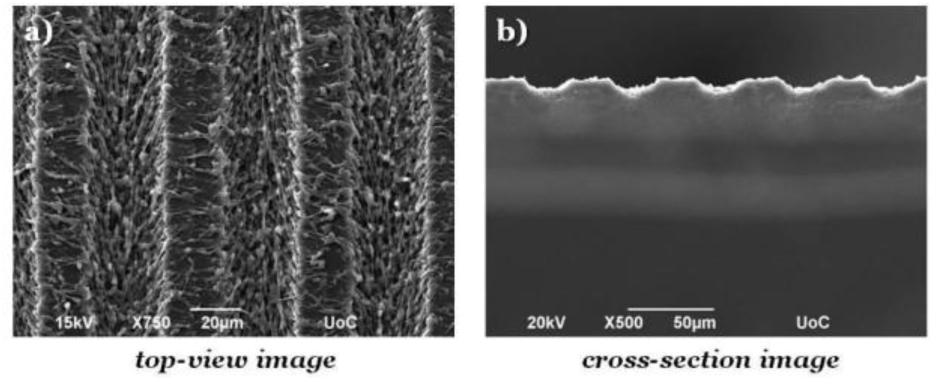
Top (a) and cross-section (b) SEM images of the polyethylene terephthalate microgrooved (PET-MG) substrates.

### 3.2 SW10 Cells’ Orientation under Static and Dynamic Culture Conditions

To examine the SW10 cells’ response on the PET-Flat and PET-MG substrates, under static and dynamic culture conditions, a series of immunocytochemical experiments were performed. Figure 3 illustrates SW10 cells cultured on both types of substrates for static and dynamic culture conditions, under the flows of 50 and 200 μL/min, respectively, on the third day of culture. We noticed, in all cases, that the SW10 cells attached strongly and proliferated well on the substrates. Additionally, cells evenly adhered and proliferated both on the top (Figure 8S in the Supplementary Material) as well as inside (Figure 3) the MG of the PET-MG substrates. The orientation of the cytoskeleton, on the top and inside the MG, has been extracted from the respective images and plotted as polar plots, depicted in Figures 9S (Supplementary Material) and 4 respectively.

**Figure 3:**
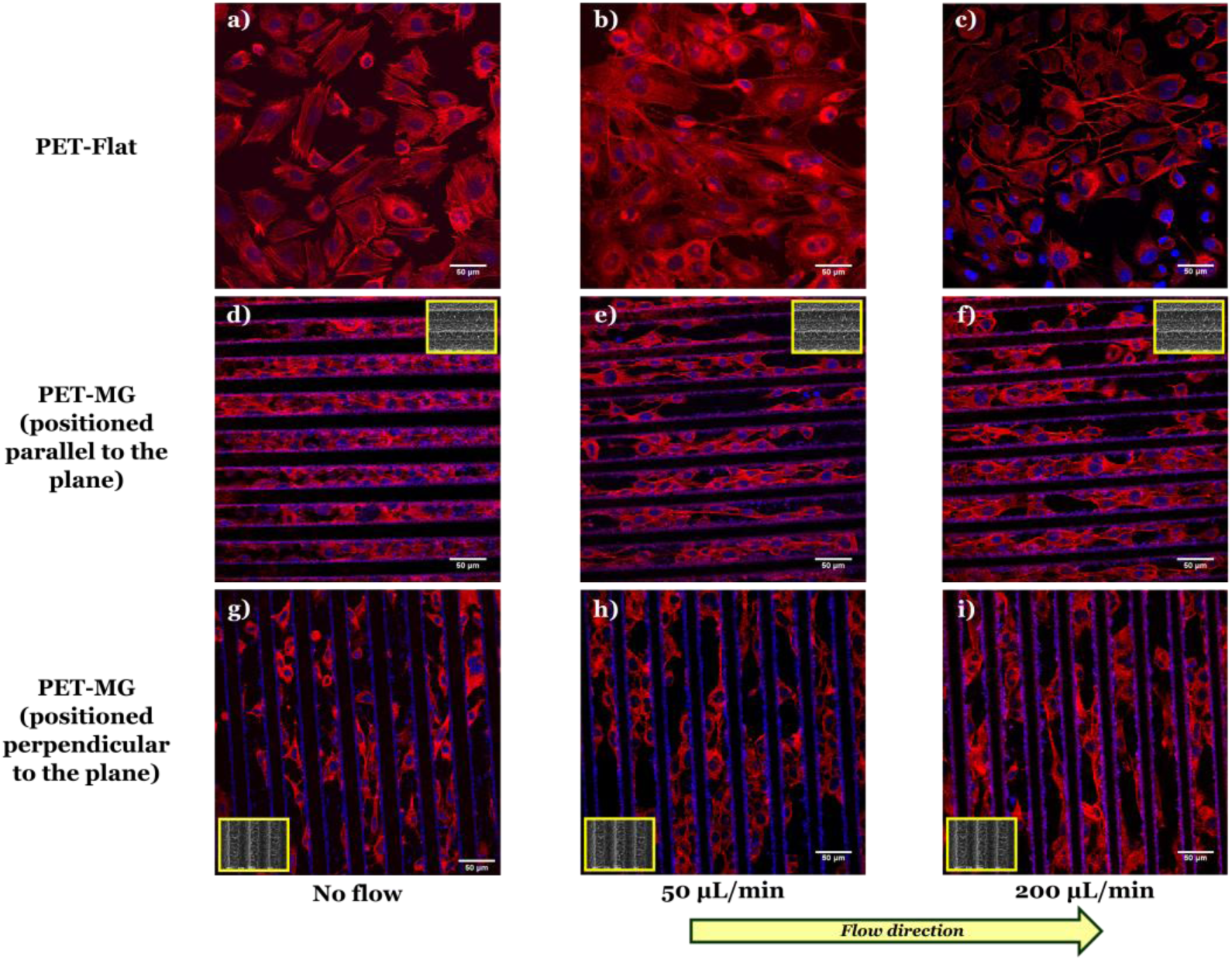
Confocal images of SW10 cells cultured on the PET-Flat (a, b, c) or inside the MG of the PET-MG substrates (d-i), under static (a, d, g) or dynamic conditions, applying 50 (b, e, h) and 200 (c, f, i) μL/min, on the third day of culture. The cytoskeleton of the cells is visualized with red color (Alexa Fluor® 680 Phalloidin) while the nuclei with blue color (DAPI). The direction of the flow was parallel (e, f) or perpendicular (h, i) to the microgrooves. The inset SEM images, framed by a yellow box, depict the geometry of microgrooves.

In particular, Figures 3a to c depict the SW10 cells cultured on the PET-Flat substrates under static and dynamic conditions. Contrary to the case without flow, we observed a preferential orientation of the cytoskeleton parallel to the flow, which is more enhanced as the flow rate is increased. This can be also clearly evidenced by the directional polar plots of cells’ cytoskeleton presented in Figure 4a. Indeed, the polar plot corresponding to the flow rate of 200 μL/min (blue line), exhibits a much narrower distribution compared to 50 μL/min (red line) at the direction of the flow, ∼ ±15°. In contrast, under static culture conditions (Figure 4a, black line) the polar plot shows a broad distribution indicating an omnidirectional cell orientation.

**Figure 4:**
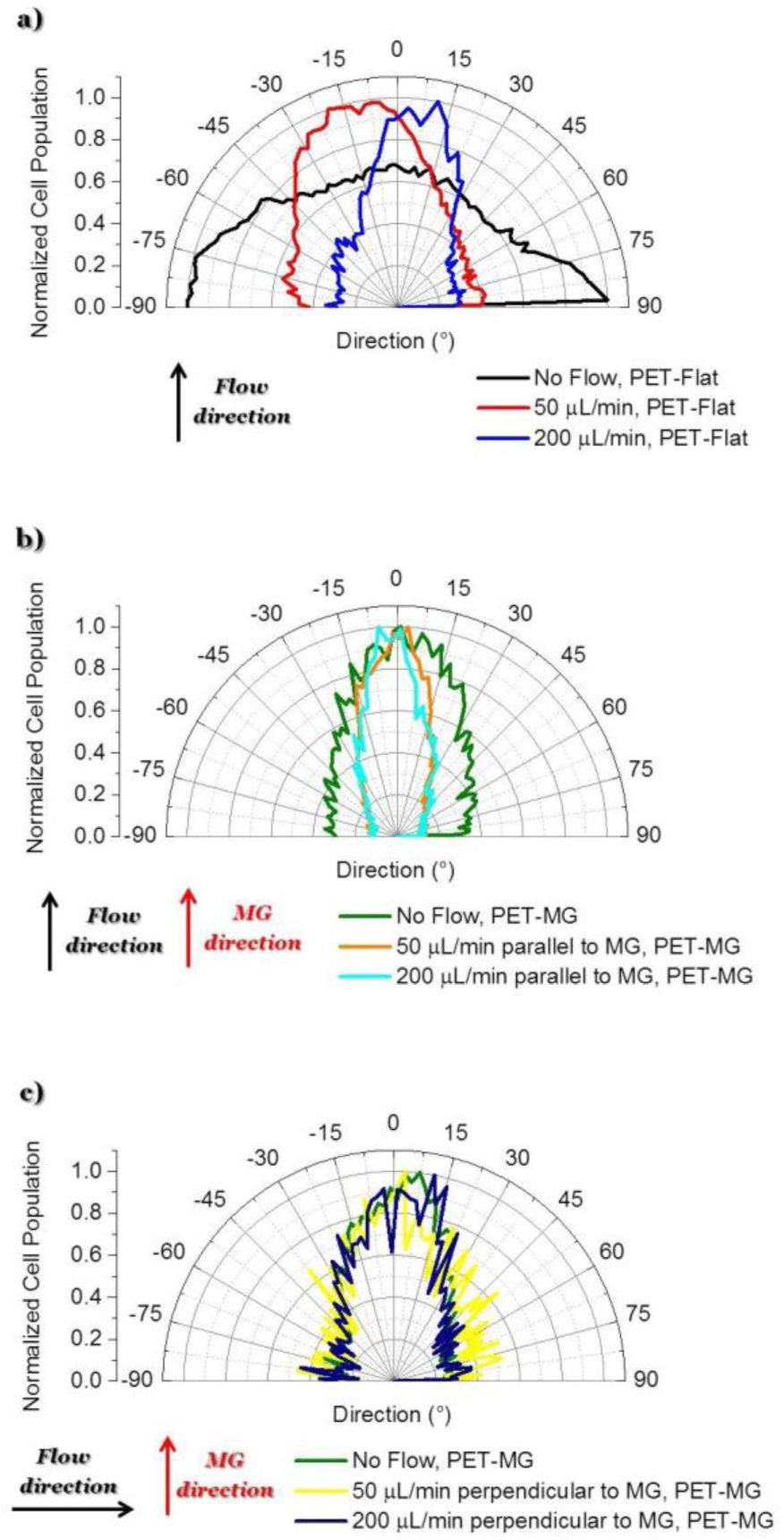
Directional polar plots of cells’ cytoskeleton on the PET-Flat or inside the MG of the PET-MG substrates. Specifically: a) No flow, PET-Flat (black line) - 50 μL/min, PET-Flat (red line) - 200 μL/min, PET-Flat (blue line), b) No flow, PET-MG (green line) - 50 μL/min parallel to MG, PET-MG (orange line) - 200 μL/min parallel to MG, PET-MG (turquoise line), c) No flow, PET-MG (green line) - 50 μL/min perpendicular to MG, PET-MG (yellow line) - 200 μL/min perpendicular to MG, PET-MG (dark blue line). The black and red arrows represent the direction of the flow and the microgrooves, respectively.The statistical analysis of the data was performed using post hoc Tukey HSD test. A p-value < 0.05 was considered significant. In particular: a) No flow, PET-Flat is significantly different from 50 μL/min, PET-Flat and 200 μL/min, PET-Flat (** p<0.01). Moreover, 50 μL/min, PET-Flat is significantly different from 200 μL/min, PET-Flat (** p<0.01); b)No flow, PET-MG is significantly different from 50 μL/min parallel to MG, PET-MG and 200 μL/min parallel to MG, PET-MG (* p<0.05). No significant difference was observed between 50 μL/min parallel to MG, PET-MG and 200 μL/min parallel to MG, PET-MG; c) No significant difference was observed between No flow, PET-MG, 50 μL/min perpendicular to MG, PET-MG and 200 μL/min perpendicular to MG, PET-MG.

Figures 3d to f show the SW10 cells seeded inside the MG of the PET-MG substrates under static and dynamic culture conditions, where the flow was applied parallel to the MG length, at the rates of 50 and 200 μL/min respectively. We observed that in both static and dynamic culture conditions, the cytoskeleton is elongated parallel to the MG length direction (Figures 3d-f). However, an enhanced cytoskeleton orientation was observed under dynamic culture conditions combined with the MG topography, as evidenced by the directional polar plots, illustrated in Figure 4b. Indeed, the normalized cell population, in this case, exhibits a narrower distribution compared to the static cultures (Figure 4b). This distribution does not seem to be affected by increasing the flow rate from 50 to 200 μL/min. Similar results on the orientation of the cytoskeleton were noticed on top of the MG of the PET-MG substrates (Figure 9Sa in the Supplementary Material).

Finally, by applying the same flow rates perpendicular to the MG length (Figures 3h-i), we observed that the cells’ cytoskeleton orientation was practically unaffected and remained elongated along the MG. This is also confirmed by the respective polar plots, presented in Figure 4c. The results were in agreement with the cytoskeleton orientation on top of the MG of the PET-MG substrates (Figure 9Sb in the Supplementary Material).

### 3.3 SW10 Cells’ Elongation under Static and Dynamic Culture Conditions

Figure 5 depicts the estimated length of SW10 cells on the PET-Flat and PET-MG substrates under static and dynamic culture conditions. We observed that the cell length, on the PET-Flat substrate, under the flow rate of 50 μL/min was significantly higher compared to the static cultures (Figure 5a). While a further elongation of the cytoskeleton was observed upon increasing the flow rate to 200 μL/min (Figure 5a). More importantly, the cell length was further enhanced upon dynamic culture conditions combined with the MG topography (Figure 5b). On the contrary, the cell length tends to decrease upon a flow perpendicular to the microgrooves (Figure 5c).

**Figure 5:**
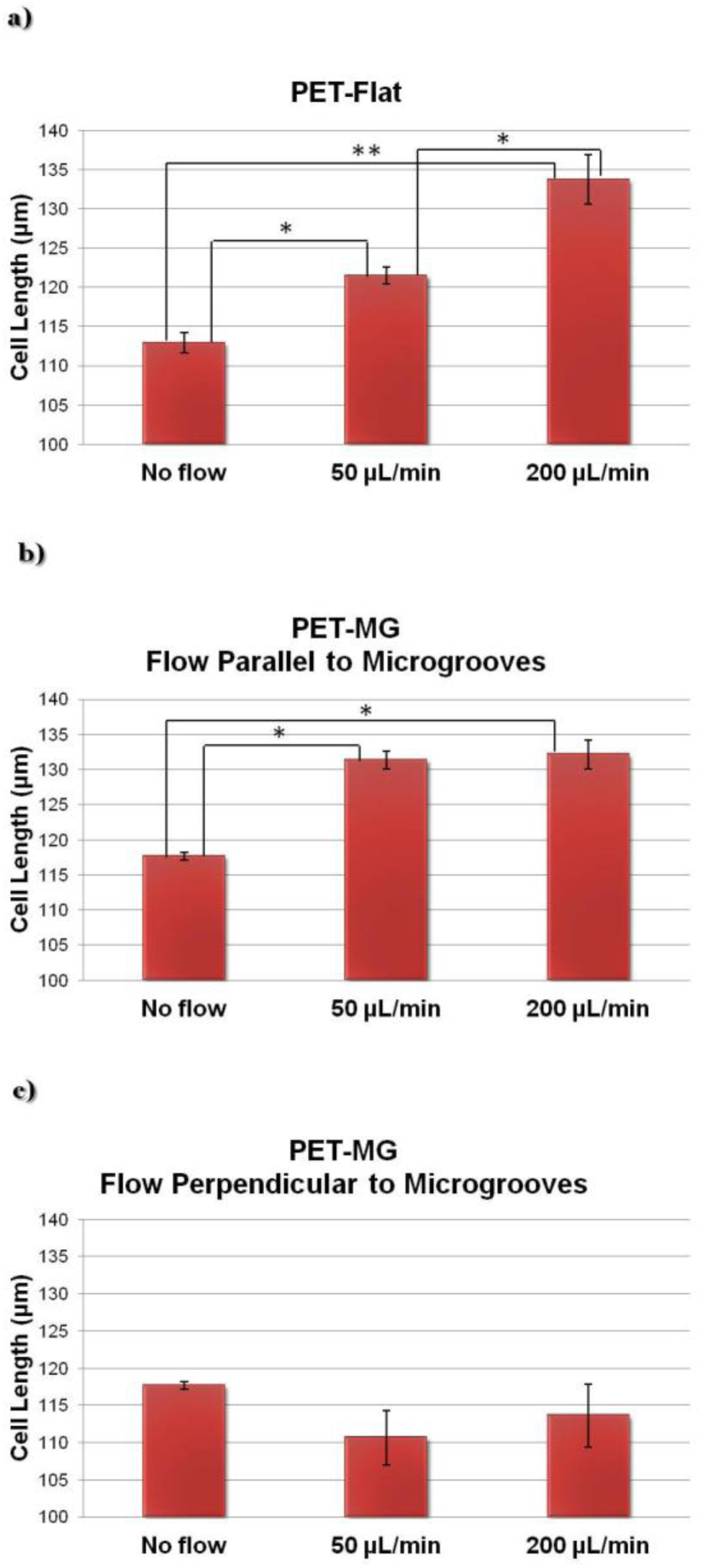
Cell length of SW10 cells (via Fiji ImageJ and Operetta High-Content Imaging System) on the PET-Flat (a) and PET-MG substrates (b, c) under static and dynamic conditions, applying 50 and 200 μL/min, on the third day of culture. The flow was parallel (b) or perpendicular (c) to the microgrooves. The statistical analysis of the data was performed using post hoc Tukey HSD test. A p-value < 0.05 was considered significant.

### 3.4 Numerically computed wall shear stress distribution on the PET-Flat and PET-MG substrates

Results from numerical simulations of the flow field in the microfluidic system chamber with steady flow at 50 and 200 μL/min flow rates for PET-Flat and PET-MG substrates are shown in Figure 6. The width and the maximum depth of the profile of MG applied to the flat substrate in the latter case was 28.68 μm and 8.87 μm respectively, as extracted from the respective SEM measurements (Figure 2). Exploiting the periodicity of the pattern of the microstructured substrate only part of the physical domain was modelled in the simulation to extract the primary features of the flow field. The simulated region, in the case where MG runs parallel to mean flow, includes a single MG and its lateral flat ridges which are assumed to be adjacent to the physical model symmetry plane. The wall shear stress distribution on the flat square cell culture substrate surface in the microfluidic system chamber indicates that the inlet and outlet triangular shaped regions of the physical model affect the velocity field in the square-shaped cell culture chamber region and reduce the area where cells are exposed to a uniform shear loading. This effect becomes more pronounced as the flow rate is increased.

**Figure 6:**
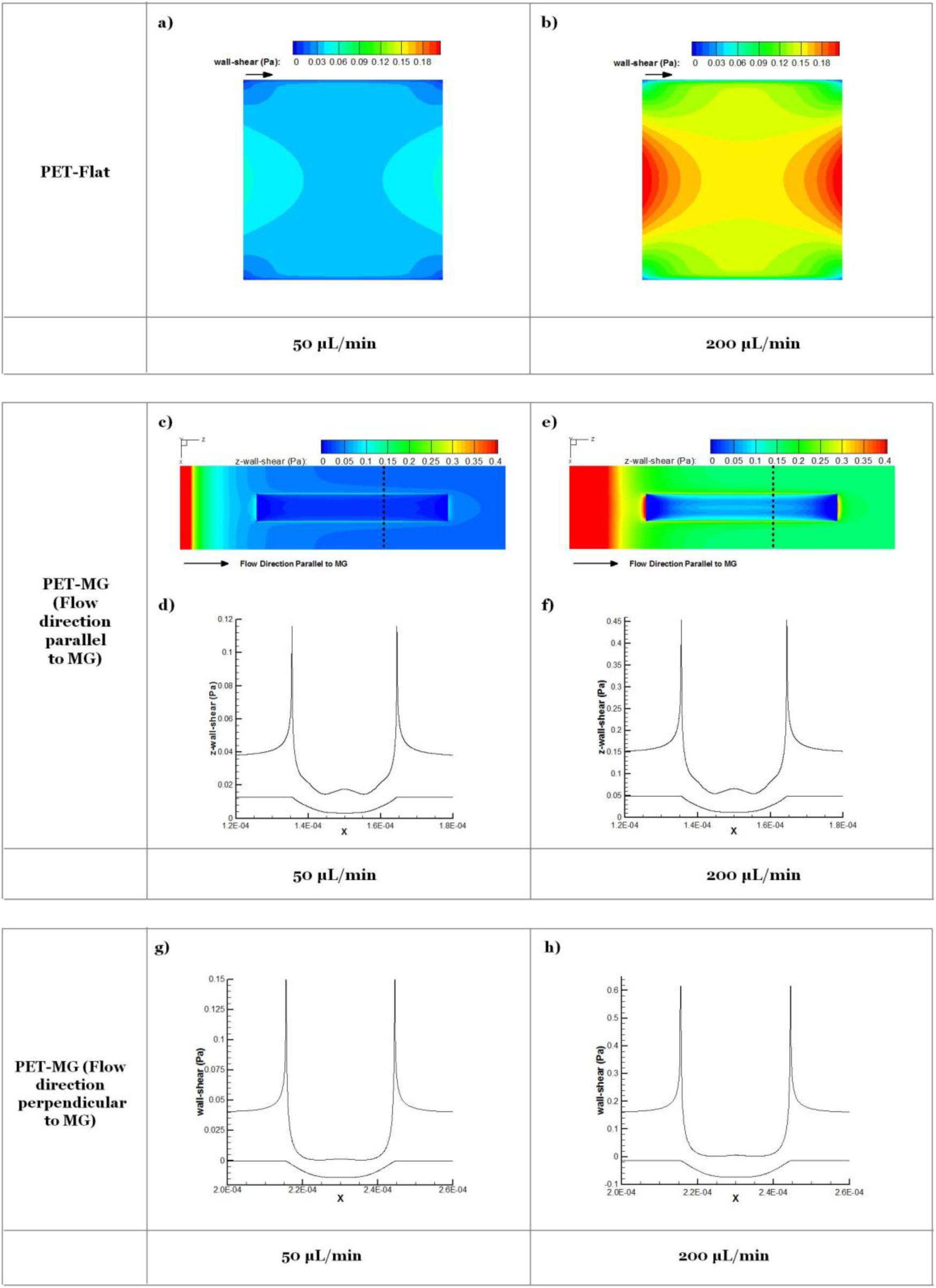
Contour plots on PET-Flat (a, b) and PET-MG substrates (c,e) and cross section profiles (dotted line) (d,f,g,h) of computed wall shear stress for 50 (a, c, d, g) and 200 μL/min (b, e, f, h) flow rates. Microgrooves are parallel (c-f) or perpendicular (g, h) to the flow direction. Lower curve in d, f, g, h depicts modelled geometry of microgroove cross section.

The addition of MG parallel to flow direction increases the complexity of the flow field which becomes three-dimensional. Cells inside the MG of the PET-MG substrates will be exposed to lower wall shear stress compared to those on the top of MG. In the case where MG are parallel to the flow, a recirculation region develops as the flow enters and exits the microgrooved channel which is characterized by relatively low wall shear stress. In the case where MG are placed perpendicular to the flow, the computational domain can be reduced to 2 dimensions exploiting the symmetries in the microfluidic system chamber and the periodicity of the MG pattern. In this case, the strong variability of wall shear stress along the MG width is evident. A strong wall shear stress gradient develops at the edges of the MG perpendicular to the flow direction. Although the flow patterns are the same in both 50 and 200 μL/min cases the gradients are as expected larger in the higher flow rate case.

## 4. Discussion

Although the axons of the PNS spontaneously regenerate after an injury, the regeneration that takes place is rarely functional because axons are usually directed in inappropriate targets [1,44,45]. Therefore, the discovery of successful methods to guide neurite outgrowth in a controllable manner, *in vitro*, is essential towards neurogenesis.

Neuronal survival and function are greatly dependent on the support of glial cells. Indeed, glial cells due to their ability to release neurotrophic factors, to express cell surface ligands and synthesize ECM, as well as, to their oriented shape and structural reorganization, provide molecular and topographical guidance stimuli for the development and outgrowth of neurons [33]. Therefore, evaluating glial cells’ response on the culture substrate topography as well as on the surrounding mechano-environment and the applied shear stress is of significant value.

Previous studies have shown that the culture substrate topography, particularly in the form of continuous electrospun polymeric fibers and grooves, as well as discontinuous isotropic and anisotropic pillars, influence neurite growth, orientation, and differentiation [7,8,10–13].

In the present study, polymeric continuous MG (PET-MG substrates) have been prepared by ultrafast laser direct writing (Figure 2) [18]. The width of the MG was chosen to be in the range of 2 to 30 μm, which is considered to be optimal for the alignment of Schwann cells [4]. Indeed, we observed that the cells’ cytoskeleton appeared to be oriented along the direction of the MG (Figures 3d, g and Figures 8Sa, d in the Supplementary Material), whereas it showed a random orientation on flat PET substrates (Figure 3a). This was also confirmed by the directional polar plots of cells’ cytoskeleton, presented in Figures 4a-b and 9Sa (Supplementary Material). Based on these results, the MG provided a favorable environment for the alignment of Schwann cells, which is crucial for axonal regeneration by providing physical guidance as well as neurotrophic and neurotropic support for axonal regrowth [33].

Apart from topography, mechanical stress is also a significant component of the host environment, as it influences the cellular signal transduction and the behavior of various cells. Indeed, previous studies have shown that fluid-induced shear stress enhances the cells’ alignment, through the reorganization of the cytoskeleton, in a variety of cell types [41,46–51]. An association between extracellular matrix alignment and cell shear stress was also shown in [52]. Thus, the fluid-induced shear stress may be also crucial for guiding neurite outgrowth. Indeed, it is known that the cell soma and the neurites of neurons are correlated to the cytoskeleton, that senses the mechanical stimuli producing different cellular responses [1,33,53]. However, the effect of shear stress on Schwann cells has been rarely reported [28,29,33]. In particular, Chafik et al. [33] reported that shear stress is a critical component of the natural environment for the regeneration of axons. They showed that mechanical stimuli (1.33 Pa for 2 hours) enhanced the proliferation of Schwann cells and caused a slight movement from their original positions. Furthermore, Gupta et al. [28,29] reported that shear stress (3.1 Pa for 2 hours) enhanced the proliferation of primary Schwann cells and down-regulated myelin protein gene expression. However, the main limitation in these studies was that the flow was not continuous, as it happens *in vivo*, but only for some hours per day. In addition to this, the effect of shear stress on the alignment of Schwann cells, as well as, the cell length has not been addressed. Finally, to the best of our knowledge, the combined effect of shear stress and topography on Schwann cells behavior has not been reported yet.

In this work, we developed a precise flow controlled microfluidic system with a custom-made geometry chamber incorporating the substrates (PET-Flat and PET-MG substrates) (Figure 1) and investigated the Schwann cells’ adhesion, orientation, and elongation under continuous flow conditions. Due to the fact that the *in vivo* values of forces during nerve regeneration are still unknown, parametric studies were performed to conclude to the flow rates of 50 and 200 μL/min (0.04 and 0.15 Pa, respectively). Higher values of shear stress caused the detachment of the cells from the culture substrates.

Under both flow rates of 50 and 200 μL/min, we observed that the cytoskeleton oriented parallel to flow, whereas it showed a random orientation under static conditions on the flat PET substrates (Figures 3a-c and 4a). Moreover, the cell length under the flow rates of 50 and 200 μL/min on the PET-Flat substrate significantly increased by 7.6% and 18.5%, respectively, compared to the static culture conditions (Figure 5a). To date, the values of shear stress reported to be stimulatory for Schwann cells for short periods are higher than 1 Pa [28,29,33]. In this study, although the values of shear stress exerted to the cells were lower, i.e. equal to 0.04 and 0.15 Pa, it is found that they were adequate to promote the cells’ alignment and elongation. Both phenomena could be attributed to the continuous fluid-induced shear stress experienced by the cells, which induces mechanical stimulation resulting in the reorganization of the cytoskeleton [1,33,53].

In a novel approach, we have studied the combined effect of shear stress and topography on Schwann cells’ behavior. Our results presented in Figures 3d-f, 8Sa-c (Supplementary Material), 4b, 9Sa (Supplementary Material) and 5b clearly show that this effect is synergetic and gives rise to further enhancement of cells’ orientation and elongation. Indeed, a significantly increased cell length by 11.7% and 12.3% was observed upon applying the flow parallel to the MG length, compared to the static culture conditions on the PET-MG substrates. However, the shear stress cannot provide further significant enhancement of cell length and orientation above 50 μL/min on MG. This indicates that small values of flow rates are sufficient to attain the maximum cell response.

When the flow was perpendicular to MG length direction, we observed that Schwann cells retained their orientation along the direction of the MG, despite the presence of shear stress in the perpendicular direction (Figures 3h-i, 8Se-f in the Supplementary Material, 4c and 9Sb in the Supplementary Material). Furthermore, the cell length, in that case, was lower compared to the static cultures on the same substrates (Figure 5c). Both results indicate that an antagonistic effect between the shear stress and topography takes place in this case and the topography effect on cell response is more pronounced.

Despite the different shear stress values on top and inside the microgrooves of the PET-MG substrates, obtained from numerical simulations, similar results on the cytoskeleton orientation were observed both on top and inside the microgrooves of the PET-MG substrates (Figures 9S in the Supplementary Material and 4b-c respectively). This indicates that the topography is the dominant factor that drives the cell orientation, compared to the shear stress (at least within the range of shear stress values used in this study.

Results obtained from numerical simulations of the flow in the cell culture chamber for PET-Flat and PET-MG substrates (Figure 6) indicated that on the flat PET shear forces are non-uniformly distributed with stronger spatial gradients located near the boundaries of the flow domain. When the substrate topography is altered by placing MG, in either parallel or perpendicular to the mean flow direction, wall shear stress distribution patterns are strongly affected with marked spatial gradients appearing at the edges of the MG profile. As the cell size is of the scale of the MG pattern it is expected that cells lying to the top of the MG will be exposed to both a high axial shear and a strong axial shear gradient in a direction perpendicular to the flow. Therefore the response of these cells regarding cytoskeleton length and orientation will be the result of a triplet of factors, i.e. a) substrate topography, b) shear stress and c) spatial gradient of shear stress. These factors appear to act in a synergistic way when MG are parallel to the mean flow direction, while the topography seems to be superior in the case where MG are placed perpendicular to the mean flow. The presence of spatial gradients of wall shear stress in the microfluidic system chamber and the effects of these gradients on cell response is of interest.

Experimental observations in cultured endothelial cells have indicated that cell morphology and phenotype response are sensitive to spatial gradients of wall shear stress [54,55]. Endothelial cells tend to align with the flow under uniform wall shear stress but appear more randomly oriented when exposed to shear stress gradients. Based on these observations it could be ascertained that shear gradients would be expected to act in a way to reduce the heights of bars in Figures 5b, c. However, as cell response shown in Figure 5 accounts for topography, magnitude, and gradients of wall shear stress, it is not possible to separate the effect of gradients. It should also be noted that as gradients of shear increase with flow rate their relative effect will be more pronounced in the 200 μL/min case. This may explain why cell length appears similar for 50 and 200 μL/min in the microstructured substrate case presented in Figure 5b in contrast to the marked length difference observed in the flat substrate case presented in Figure 5a.

It should be noted that the computational geometry of the PET-MG substrate is an idealized version of the real microstructured groove geometry depicted in Figure 2. Therefore although the pattern in the wall shear stress gradients will be similar minor quantitative differences in the experimental conditions are expected. This, however, is not expected to alter our main findings.

## 5. Conclusions

The combined effect of shear stress and topography on SW10 cells’ adhesion, orientation, and elongation has been investigated under both static and dynamic culture conditions. The experimental findings were analyzed together with numerical simulations of the distribution of flow-induced wall shear stress which is strongly affected by the substrate topography. It is revealed that, depending on the relation of the direction of flow with respect to the topographical features, wall shear stress gradients may be acting in a synergistic or antagonistic manner to topography in promoting guided morphologic cell response. Our results demonstrate the ability to guide the outgrowth of Schwann cells *in vitro* that could be potentially useful in the fields of neural tissue engineering with the creation of autologous graft substitutes for nerve tissue regeneration.

## 6. Acknowledgments

This work was supported by NFFA (EU H2020 framework programme) under grant agreement n. 654360 from 1/9/2015 to 31/8/2019; State Scholarship Foundation (IKY) within the framework of the Action “Doctoral Research Support” (MIS 5000432), ESPA 2014-2020 Program, CN: 2016-ESPA-050-0502-5321; Onassis Foundation through the G ZM 039-1/2016-2017 scholarship grant; and European Horizon 2020 - H2020-EU.1.2.1. - FET Open Program under Grant No. 829060 “NeuroStimSpinal”. We acknowledge also support of this work by the project HELLAS-CH (MIS 5002735) implemented under “Action for Strengthening Research and Innovation Infrastructures”, funded by the Operational Programme “Competitiveness, Entrepreneurship and Innovation” (NSRF 2014-2020) and co-financed by Greece and the European Union (European Regional Development Fund).

## Notes

### Competing Interest Statement

The authors have declared no competing interest.

